# The histone chaperone FACT modulates nucleosome structure by tethering its components

**DOI:** 10.1101/309708

**Authors:** Tao Wang, Yang Liu, Garrett Edwards, Daniel Krzizike, Hataichanok Scherman, Karolin Luger

## Abstract

Human FACT (hFACT) is a conserved histone chaperone that was originally described as a transcription elongation factor with potential nucleosome assembly functions. Here we show that FACT facilitates tetrasome assembly and H2A-H2B deposition to form hexasomes and nucleosomes. In the process, FACT tethers components of the nucleosome through interactions with H2A-H2B, resulting in a defined intermediate complex comprised of FACT, a histone hexamer and DNA. Free DNA extending from the tetrasome then competes FACT off H2A-H2B, thereby promoting hexasome and nucleosome formation. Our studies provide mechanistic insight into how FACT may stabilize partial nucleosome structures during transcription or nucleosome assembly, seemingly facilitating nucleosome disassembly and nucleosome assembly.

## INTRODUCTION

The organization of all genomic DNA into nucleosomes represents a formidable barrier to the cellular machinery acting on DNA. The tight wrapping of 147 base pairs (bp) of DNA around a histone octamer prevents access of DNA and RNA polymerases and of regulatory factors (Fischle et al., 2003, Groth et al., 2007, Kulaeva et al., 2007, Li et al., 2007, Price et al., 2013). Therefore, the mechanism by which nucleosomes are altered dynamically during these processes is the topic of intense studies.

Nucleosomes are modular complexes whose assembly begins with the deposition of one (H3-H4)_2_ tetramer onto ~70 bp of DNA to form a ‘tetrasome’ (Mattiroli et al., 2017), followed by the addition of two H2A-H2B dimers that are stabilized through multiple interactions with the (H3-H4)_2_ tetramer and flanking 2×35 bp of DNA (Luger et al., 1998). Nucleosome disassembly likely occurs through the reverse pathway. Outside of the nucleosome, the highly basic histones are found in complex with histone chaperones that prevent their nonspecific interactions with DNA and orchestrate their sequential deposition onto DNA (Das et al., 2010, De Koning et al., 2007, Eitoku et al., 2008).

FACT (<u>FA</u>cilitates <u>C</u>hromatin <u>T</u>ranscription) is an abundant and essential histone chaperone that is conserved in all eukaryotes. Human FACT (hFACT) is a heterodimer composed of Spt16 and SSRP1 (Orphanides et al., 1999). Yeast FACT additionally requires Nhp6, an HMG-B domain subunit that in metazoans is fused to SSRP1 (McCullough et al., 2018). The FACT complex interacts with all three RNA polymerases (Birch et al., 2009, Tessarz et al., 2014) and facilitates transcription by disrupting nucleosomes in their path, and by aiding in the re-deposition of histones post-transcription (Belotserkovskaya et al., 2003, Formosa et al., 2002).

Electrophoretic mobility shift assays (EMSA) suggest that yeast FACT binds to H2A-H2B dimer and (H3-H4)_2_ tetramer with similar affinity (Kemble et al., 2015, Kemble et al., 2013). Interactions between yeast FACT and H2A-H2B dimers are promoted through short acidic regions near the C-termini of each subunit that bind H2A-H2B dimer competitively and preclude H2A-H2B binding to DNA (Kemble et al., 2015, Kemble et al., 2013). The interaction between FACT and the (H3-H4)_2_ tetramer was confirmed in the structure of a portion of human Spt16 with a (H3-H4)_2_ tetramer. This structure suggests that the FACT-(H3-H4)_2_ tetramer complex is incompatible with the interactions of H3-H4 with DNA within the nucleosome (Tsunaka et al., 2016). Since H2A-H2B and H3-H4 bind distinct regions on FACT, they can bind to FACT simultaneously in the absence of DNA (Tsunaka et al., 2016).

FACT was originally described as a transcription enlongation factor (Orphanides et al., 1998), but how FACT affects nucleosome structure is unknown. Using a defined chromatin template, it was shown that FACT displaces one or two H2A-H2B dimers from nucleosomes during transcription (Belotserkovskaya et al., 2003, Hsieh et al., 2013). FACT interaction with H2A-H2B is essential for this activity. ChIP experiments in yeast suggest that yFACT also reassembles nucleosomes in the wake of RNA polymerase II (Jamai et al., 2009, Nguyen et al., 2013). Incorporation of new H3 in yeast gene bodies increases in the absence of Spt16 (Voth et al., 2014), suggesting that FACT contributes to the maintenance of pre-existing tetrasomes. Thus, FACT appears to act both as a nucleosome assembly and disassembly factor. To reconcile these seemingly opposing functions, it was proposed that FACT holds the components of the nucleosome in a ternary complex during transcription elongation (Formosa, 2012). Consistent with this idea, a recent study showed that hFACT does not bind to intact nucleosomes, but only to nucleosomes with a destabilized H2A-H2B dimer (obtained by reconstitution of nucleosome with two DNA fragments (33/112 bp) rather than with 147 bp DNA) (Tsunaka et al., 2016).

Here, we show that FACT interaction with H2A-H2B facilitates its binding to a (H3-H4)_2_ tetramer to form a defined ternary complex. In the presence of DNA, a FACT●(H2A-H2B complex interacts with DNA-bound (H3-H4)_2_ tetramer, thereby tethering the components of the nucleosome. Ultimately, DNA competes FACT off the H2A-H2B dimer, resulting in the formation of hexasomes and nucleosomes.

## RESULTS

### FACT forms a ternary complex with histones H2A-H2B and (H3-H4)_2_

The previously reported stoichiometry of the hFACT:H2A-H2B complex was confirmed by Sedimentation Velocity Analytical Ultracentrifugation (SV-AUC). The sedimentation coefficient of FACT is 7.3S (represented as Svedberg units (S)), corrected for standard conditions of water at 20 Celsius; Figure 1A), corresponding to a molecular weight of 198 kDa, and within error of the theoretical molecular weight (Table 1). Addition of equimolar amounts of H2A-H2B (which itself sediments with an S-value of 1.7 to 2.5S, Figure 1B) increases the rate of sedimentation to 8.3S, and the resulting apparent molecular weight is consistent with a FACT to H2A-H2B stoichiometry of 1:1 (Table 1). Doubling the amount of H2A-H2B does not further increase the rate of sedimentation (Figure 1A). The distribution of S-values indicates a high degree of homogeneity of the FACT-histone complex.

**Figure 1.**
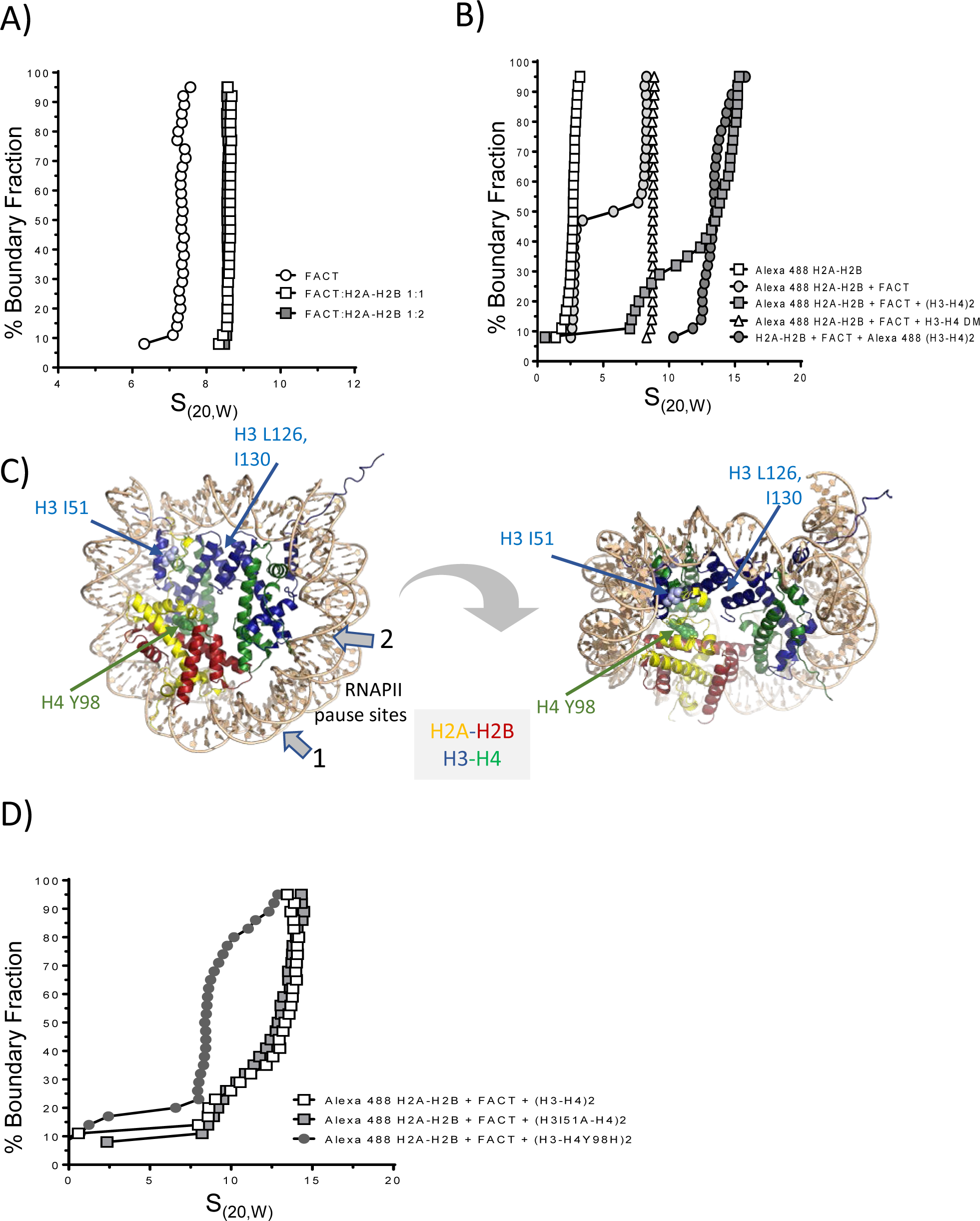
H2A-H2B facilitates FACT interaction with the (H3-H4)^2^ tetramer to form a ternary complex. (A) SV-AUC of histone-FACT complexes. Sample composition is represented in a van Holde-Weischet plot. 1.8 µM FACT was mixed with H2A-H2B at the indicated molar ratios. H2A-H2B sediments between 1.7 and 2.5S (Fig. 1B). (B) SV-AUC of histone-FACT complexes, by monitoring the sedimentation of fluorescently labeled histone H2B or H3 (*) with a fluorescence detection system (FDS; 488 nm). Open squares: 100 nM H2A-H2B*; open circles: 100 nM H2A-H2B* + 200 nM FACT; closed squares: 100 nM H2A-H2B* + 400 nM FACT + 400 nM H3-H4; closed circles: 100 nM H2A-H2B + 400 nM FACT + 400 nM H3-H4*. Open triangles: 100 nM H2-H2B*, 400 nM FACT, and 400 nM DM-H3-H4. (C) H3I51 and H4Y98 are shown in space filling representation in 1AOI. RNAPII pause sites specific to FACT are indicated with grey arrows; the location of H3L126 and I130 is also indicated. (D) SV-AUC of H2A-H2B* (100 nM) with either 400 nM H3-H4, H3I51A-H4, or H3-H4Y98H, and 400 nM FACT. Alexa 488 fluorescence was monitored with the FDS. S-values from all SV-AUC experiments are listed in Table 1

**Table 1:**
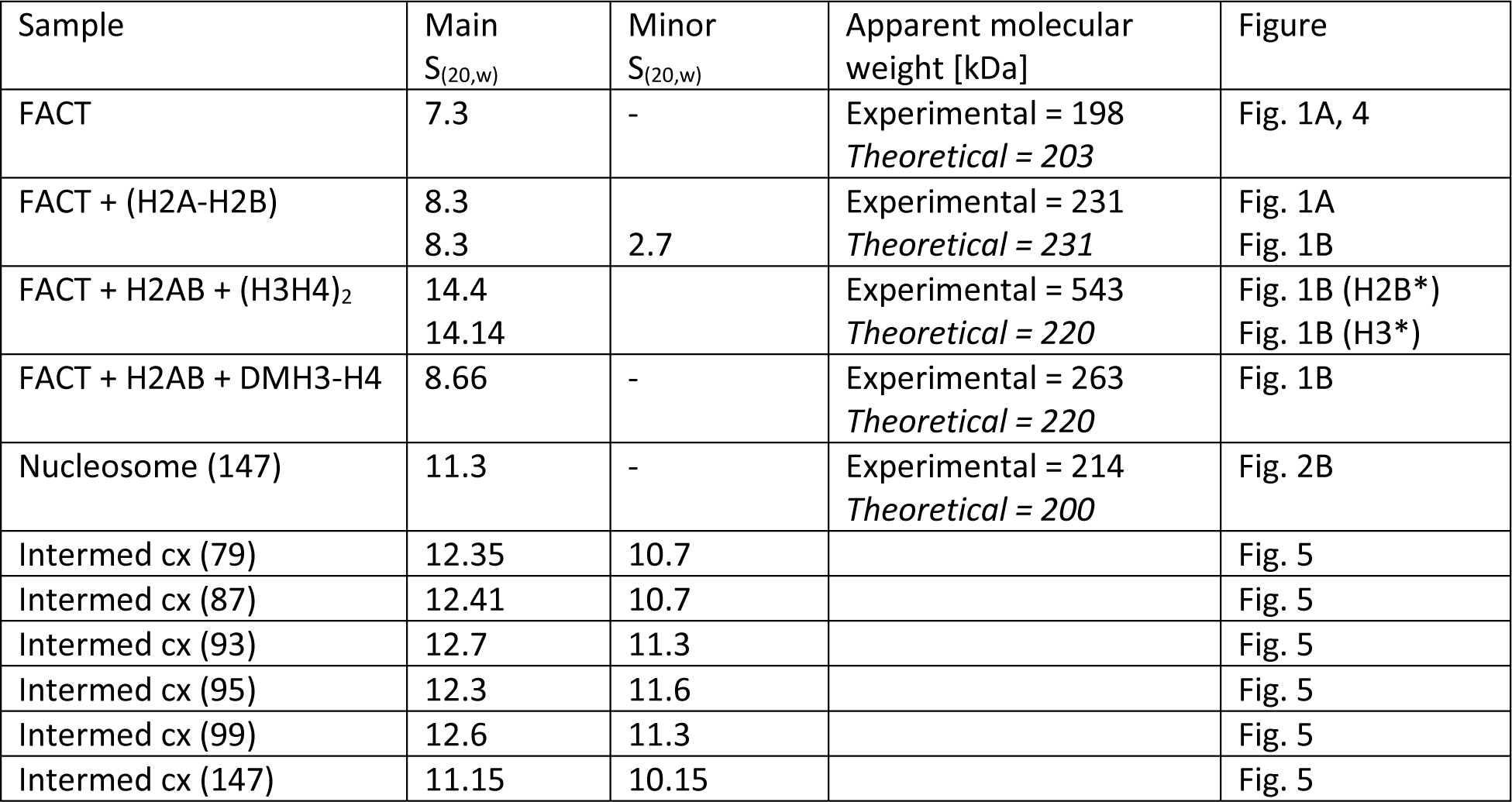
S-values and calculated molecular weights of complexes.

We attempted to use the same approach to determine the stoichiometry of the FACT-(H3-H4)_2_ tetramer complex. When 1.8 µM (H3-H4)_2_ tetramer was mixed with an equimolar amount of FACT, visible aggregates formed and only a small percentage of the input absorbance was monitored during SV-AUC. The acidic domain in Spt16 CTD interacts with H3-H4 nonspecifically (Tsunaka et al., 2016), and this may cause precipitation under SV-AUC conditions.

### hFACT binds to H2A-H2B and H3-H4 simultaneously

H2A-H2B binds to a conserved peptide motif in the yeast Spt16 CTD, whereas the (H3-H4)_2_ tetramer interacts with the hSpt16 middle domain (Kemble et al., 2015, Tsunaka et al., 2016). We confirmed the simultaneous interaction of FACT with H2A-H2B and H3-H4 (shown by (Tsunaka et al., 2016) by analytical ultracentrifugation equipped with a Fluorescence Detection System (AUC-FDS), which allows us to specifically monitor the sedimentation of fluorescently labeled constituents in complex mixtures. 100 nM Alexa 488 labeled H2A-H2B was spun in the absence or presence of 200 nM FACT. The sedimentation coefficient of H2A-H2B increases to ~8.3S in the presence of FACT (Figure 1B, Table 1), consistent with what is seen at higher concentrations (Figure 1A). Upon addition of unlabeled H3-H4 to this complex, the sedimentation coefficient distribution indicates the presence of two species, one of which can be attributed to the FACT●(H2A-H2B) complex (~8.3S), while the second species (14S) likely represents a complex of FACT, H2A-H2B, and H3-H4. To confirm this, we repeated the experiment with fluorescently labeled H3-H4 (and unlabeled H2A-H2B; FACT), and this complex had a sedimentation coefficient of ~13.8S (Figure 1B). This demonstrates that FACT binds to H2A-H2B and H3-H4 simultaneously. The apparent molecular weight calculated from these experiments is over twice the molecular weight of a FACT●H2A-H2B●(H3-H4)_2_ complex, and the stoichiometry of this assembly remains undetermined. As observed to a greater extent with unlabeled H3-H4, mixing fluorescently labeled H3-H4 (400 nM) with 200 nM FACT in the absence of H2A-H2B resulted in aggregation, as judged by the loss of > 50 % of fluorescence intensity (Figure S1). These results indicate that H2A-H2B dimer facilitates the proper interaction of FACT with H3-H4.

In these experiments, the H3-H4 concentration was kept below 500 nM, and thus H3-H4 exists in an equilibrium of H3-H4 dimer and (H3-H4)_2_ tetramer (Liu et al., 2012). To resolve whether FACT●H2A-H2B binds to an H3-H4 dimer or an (H3-H4)_2_ tetramer, we tested a mutated version of H3 that precludes tetramer formation (DMH3; L126A, I130A, C110E; Figure 1C; (Mattiroli et al., 2017). Available structural data show that these amino acids are not located in the FACT●(H3-H4)_2_ interface (Tsunaka et al., 2016), and are thus unlikely to contribute to the interaction with FACT. The addition of H3-H4 DM results in a minor shift in sedimentation of the FACT●H2A-H2B complex (Figure 1B). The molecular weight derived from this experiment indicates a FACT●H2A-H2B complex, possibly with a weakly associated single H3-H4 dimer (Table 1).

### Direct interaction of H2A-H2B with H3-H4 is essential for ternary complex formation

Next, we asked whether H2A-H2B, when in complex with FACT, directly interacts with the (H3-H4)_2_ tetramer through interactions resembling those in the nucleosome (Luger et al., 1997). To this end, we tested mutated histones H3I51A and H4Y98H that assemble into (H3-H4)_2_ tetramers and nucleosomes *in vitro* (Hsieh et al., 2013), but cannot be refolded into histone octamers (Ferreira et al., 2007); (Hsieh et al., 2013); (Ramachandran et al., 2011); Figure 1C). The replacement of H4Y98 with glycine in yeast is lethal (Santisteban et al., 1997). Neither side chain appears to be involved in the interaction with FACT (Tsunaka et al., 2016). AUC-FDS was repeated with tetramers refolded with mutant H3 or H4. We found that H3I51A does not significantly affect FACT interactions with a histone hexamer, while the more disruptive H4Y98H almost completely abolishes the integration of H3-H4 into the FACT●(H2A-H2B) complex (Figure 1D). This indicates that the interaction between the H2A docking domain and H4 is required, while the close packing of H3 αN with H2A and the histone fold of H3 is not (Figure 1C). Because H2A-H2B dimers and (H3-H4)_2_ tetramers do not interact with each other at 150 mM NaCl in the absence of DNA, our results suggest that their interaction is stabilized by FACT.

### FACT neither disassembles nor interacts with intact nucleosomes in vitro

Increasing amounts of FACT were mixed with nucleosomes that had been reconstituted with a 147 bp 601 DNA fragment. Reaction products were analyzed by 5% native PAGE, and visualized through SYBR gold staining and H2B fluorescence (Figure 2A). The intensity of the nucleosome band remained constant as FACT was titrated, indicating that under these conditions, FACT neither binds to nor disassembles nucleosomes. To exclude that this is due to the unique properties of the 601 DNA sequence, we performed the same assay with nucleosomes reconstituted with the less stable 147 bp α-satellite DNA, with identical results (Figure S2). The inability of FACT to bind fully assembled nucleosomes was confirmed by AUC. The sedimentation coefficient of a mono-nucleosome is ~11S (Yang et al., 2011) and remains unchanged upon addition of a four-fold excess of FACT (Figure 2B, Table 1). The mono-disperse sedimentation coefficient distribution indicates that no significant amounts of sub-nucleosomal complexes were formed.

**Figure 2.**
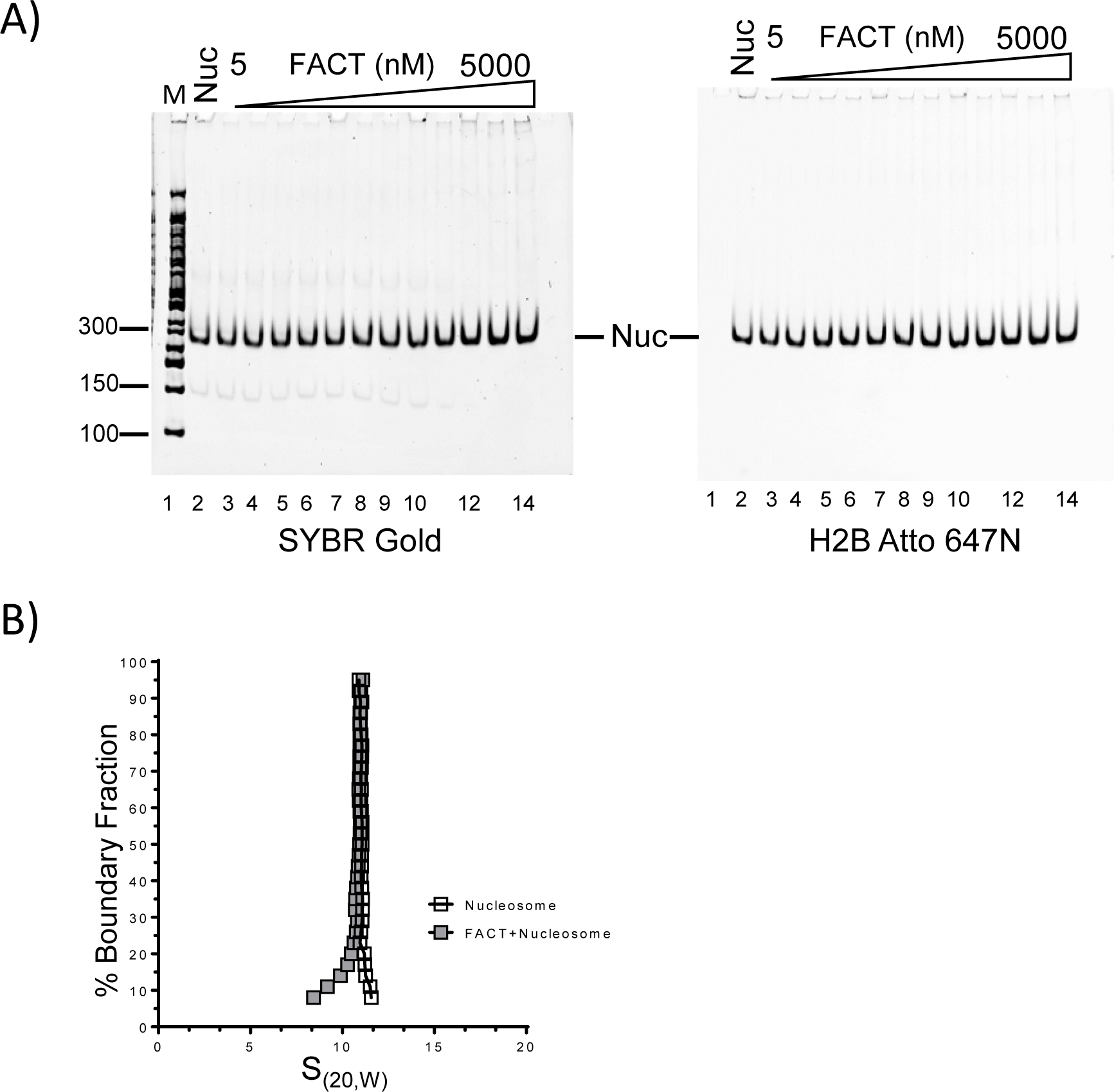
FACT neither disassembles nor binds properly assembled nucleosomes. (A) 10 nM nucleosome, reconstituted with labeled H2B on 146 bp of DNA, was incubated with increasing amounts of FACT (10 nM to 5 µM), and analyzed by native PAGE. The gel was visualized by SYBR Gold for DNA (left panel), and by H2B fluorescence (right panel). (B) The sedimentation behavior of nucleosomes remains unchanged upon addition of FACT. 130 nM nucleosomes were mixed with 520 nM FACT, and SV-AUC was performed by monitoring absorbance from DNA. Addition of FACT does not increase the S(_20,W_) value of the nucleosome.

### FACT functions as a tetrasome assembly factor, and also facilitates H2A-H2B deposition onto tetrasomes and hexasomes

We next investigated whether FACT could assemble nucleosomes. First, we tested the ability of FACT to promote the first step of nucleosome assembly, i.e. the deposition of a (H3-H4)_2_ tetramer onto DNA to form tetrasomes. FACT was preincubated with H3-H4 before adding a 147 bp 601 DNA fragment. The reaction products were analyzed by 5% native PAGE (Figure 3A). Under these conditions (15-240 nM FACT, 30 nM H3-H4), FACT does not form perceptible aggregates with H3-H4. In the absence of FACT, only a small amount of tetrasome is formed (Figure 3A, lanes 3 and 9), while increasing amounts of tetrasome appeared upon titration of FACT (Figure 3A, lanes 4-8 and 10-14).

**Figure 3.**
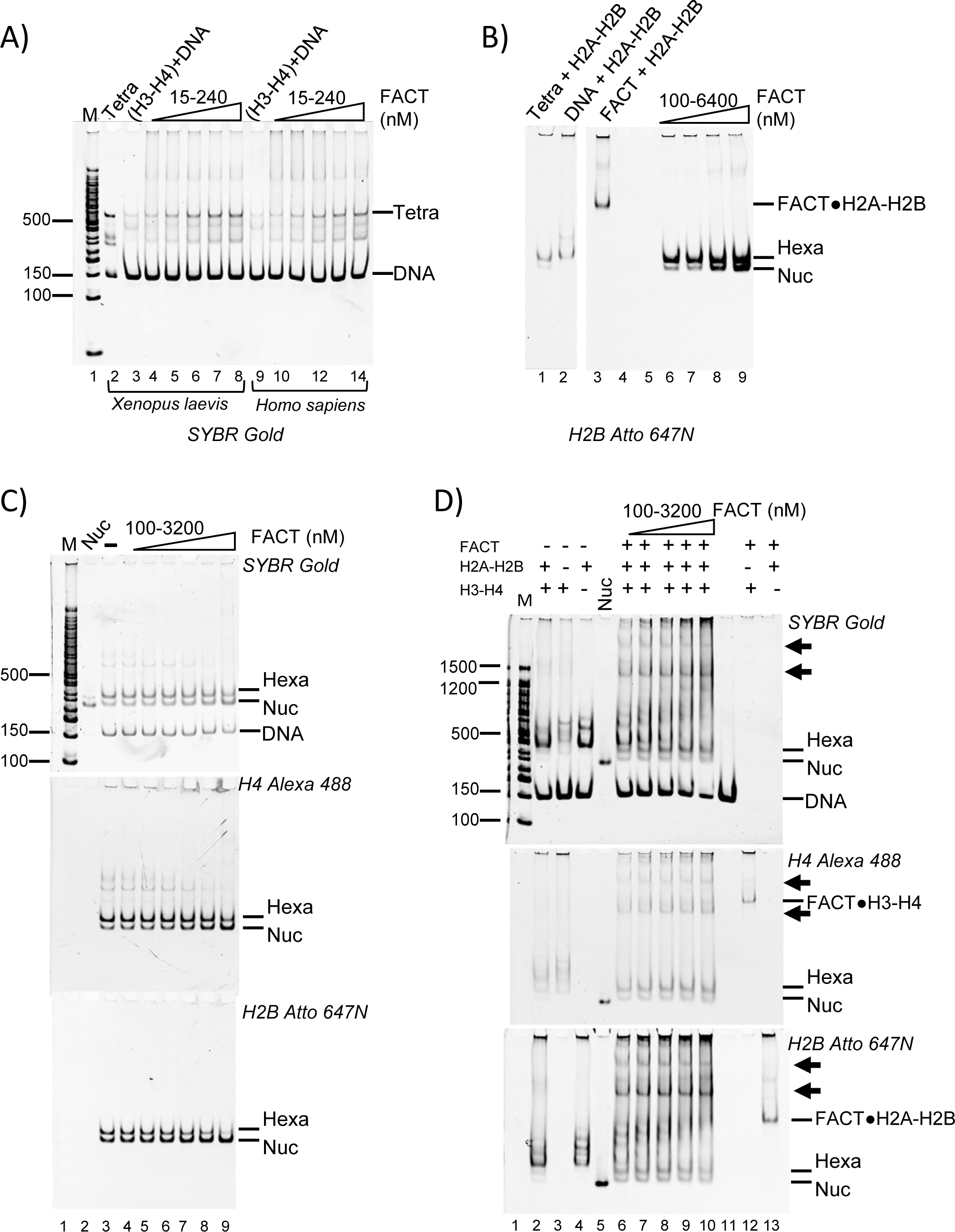
FACT has moderate nucleosome assembly activity. (A) FACT facilitates tetrasome assembly. 147-bp 601 DNA (30 nM) was incubated with FACT (15-240 nM) pre-equilibrated with 30 nM *Xenopus laevis* or *Homo sapiens* (H3-H4)_2_, separated by PAGE, and visualized by SYBR Gold (DNA). Lane 2 shows a salt-assembled tetrasome on the same DNA (Tetra). (B) FACT facilitates the deposition of H2A-H2B dimers onto the tetrasome, resulting in hexasome and nucleosome assembly. 60 nM salt-reconstituted tetrasome (lane 2) and 30 nM Atto 647N labeled H2A-H2B were incubated with increasing amounts of FACT (10-1280 nM), and gels were visualized by H2B fluorescence. (C) FACT facilitates the deposition of H2A-H2B onto the hexasome. 20 nM under-assembled nucleosome with fluorescence labels on H4 (Alexa 488) and H2B (Atto 647N) (lane 3) was incubated with increasing amounts of FACT, and the gel was visualized as indicated. (D) FACT (100 nM to 3200 nM) promotes hexasome and nucleosome assembly from free histones (50 nM H3-H4; 100 nM H2A-H2B) and DNA (25 nM). FACT (at the indicated concentrations) was incubated with histones at RT for 10 min, and then DNA was added and products were visualized as indicated. The two intermediate complexes are indicated by arrows.

We next asked if FACT facilitates H2A-H2B addition to tetrasomes that had been preassembled from 147-bp ‘601’ DNA and H3-H4 by salt dilution. Fluorescently labeled H2A-H2B was pre-mixed with the indicated amounts of FACT and incubated with tetrasomes, then analyzed by native PAGE (Figure 3B). In the absence of FACT, some nucleosome and hexasome bands are observed (Figure 3B, lane 1), but the intensity of hexasomal and nucleosomal band is increased as FACT is titrated (Figure 3B, lanes 6-9). However, a large excess of FACT is required for this effect, and hexasome is the main product under all conditions. This suggests that FACT promotes the first steps of nucleosome assembly, but is inefficient in adding the final H2A-H2B dimer to a hexasome.

To further investigate this activity, we prepared deliberately under-assembled nucleosomes by combining optimal amounts of DNA and H3-H4 with insufficient amounts of H2A-H2B in salt reconstitution. This results in a mixture of nucleosomes, hexasomes, tetrasomes, free histones and DNA after salt dialysis (Figure 3C, lane 3). As FACT is titrated, the fluorescence intensity from nucleosomal H4 increases, while the intensity of tetrasomal H4 decreases (Figure 3C; quantification in S3). Again, this requires very high FACT concentrations (i.e. a 64-fold excess of FACT over input DNA), suggesting that the addition of the final dimer to a hexasome is not a preferred reaction for FACT.

To test whether FACT assembles nucleosomes *de novo* from refolded histones and DNA, increasing amounts of FACT were mixed with a constant amount of 488-labeled H3-H4 and 647-labelled H2A-H2B before adding DNA. In the absence of FACT, histones bind nonspecifically to DNA, and very little nucleosome is formed (Figure 3D, lanes 2-4). In the presence of FACT, some nucleosomes are assembled (Figure 3D, lanes 6-10), but hexasomes are the main products, as observed before (Figure 3B). Additionally, two complexes with much higher electrophoretic mobility appear in the presence of FACT (indicated by arrows, discussed below).

To confirm the quality of the nucleosomes assembled by FACT, we compared their resistance towards micrococcal nuclease (MNase) to that of salt-assembled nucleosomes. Free FACT and intermediate complexes containing flag-tagged FACT (i.e. those indicated by asterisks in Figure 3D), were removed by anti-FLAG affinity purification, and the flow through containing nucleosome assembly products was analyzed by MNase digestion (Figure S4). Nucleosomes assembled by FACT display an MNase digestion pattern that is similar to what is observed for control nucleosomes, and unlike the pattern obtained from histones added to DNA in the absence of FACT (Table S1). Together, our data suggest that FACT has moderate nucleosome assembly activity. This is primarily through facilitating tetrasome assembly, but also through aiding H2A-H2B deposition onto tetrasomes, and, to a more limited extent, onto hexasomes.

### FACT●H2A-H2B and DNA-bound H3-H4 form a ternary complex

The high-molecular weight complexes observed by native PAGE during FACT-mediated nucleosome assembly assays minimally contain H4, H2B and DNA (Figure 3D). We speculated that these complexes form as FACT deposits H2A-H2B onto tetrasomes. To test this, we reconstituted tetrasome with 147 bp DNA and H3-H4 by salt dilution, and this was added to FACT pre-mixed with H2A-H2B (Figure 4A). FACT does not bind to tetrasome in the absence of H2A-H2B (Figure 4A, left panel, lane 4). However, when pre-bound with H2A-H2B, FACT binds to tetrasomes efficiently, and forms two intermediate complexes (Figure 4A, lanes 5-6), as observed in Figure 3D. To confirm that the complexes contain all histones, DNA and FACT, we analyzed these samples by immobilizing flag-tagged FACT and associated proteins on an affinity column, and visualized DNA, H4 and H2B fluorescence of bound and unbound fractions (Figure 4B). Nucleosomes (yellow), sub-nucleosomal complexes (green and red), and free DNA were enriched in the flow through (lane 3), while only the intermediate complexes containing DNA, H2B, and H4, eluted with the FLAG peptide (yellow, lane 4).

**Figure 4.**
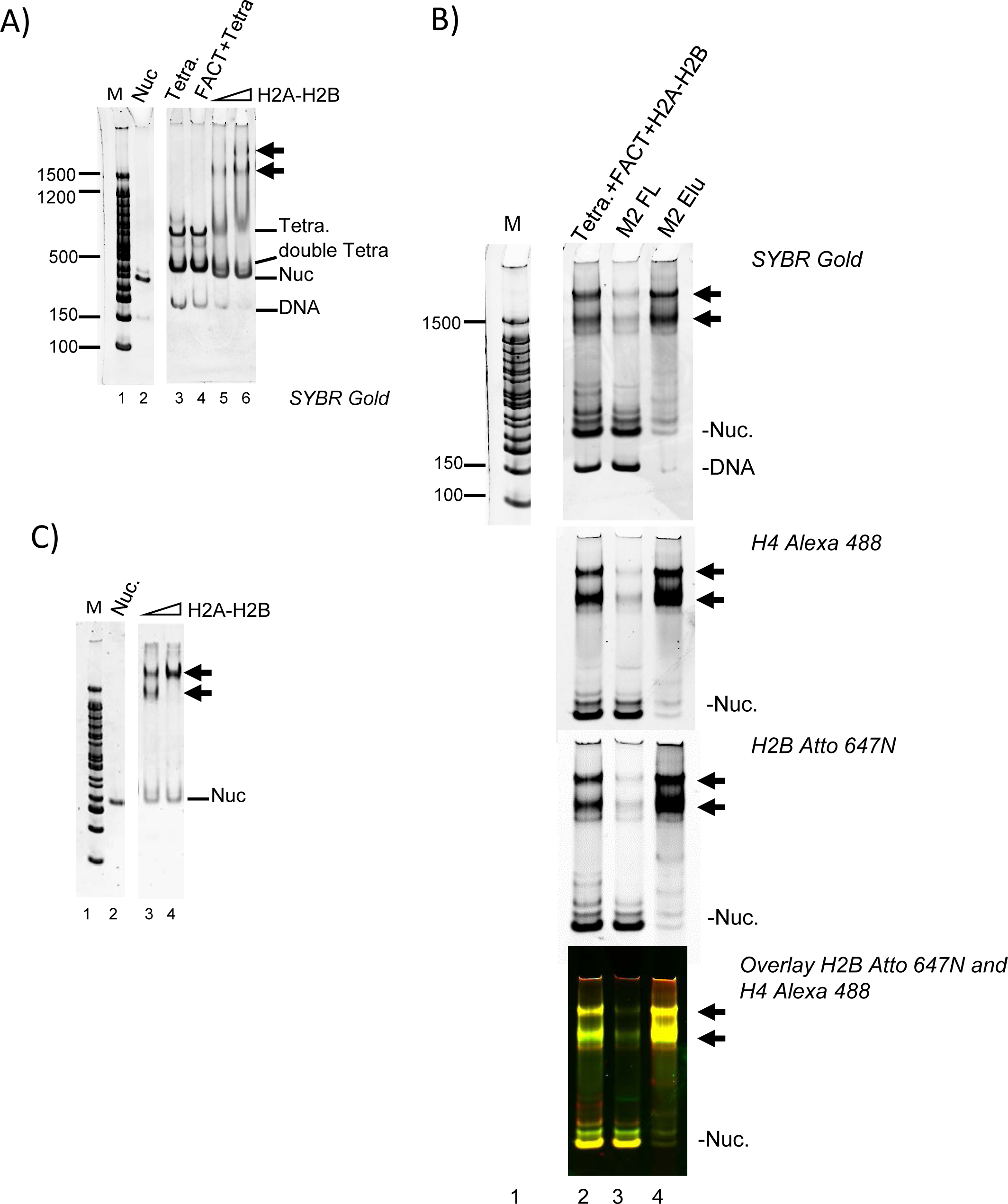
H2A-H2B promotes FACT interaction with the tetrasome. (A) FACT (800 nM) was mixed with 60 nM or 120 nM H2A-H2B before addition of tetrasome (40 nM) assembled with 147 bp DNA. Double tetra: two (H3-H4)_2_ assembled on 147 bp DNA. (B) FACT (1.3 µM) was pre-mixed with 650 nM H2A-H2B (labeled with Atto 647), and 150 nM tetrasome (labeled with Alexa 488) was added. Intermediate complexes (indicated by arrows) were enriched over M2 resin and analyzed by 5% PAGE. The marker (lane 1) is on the same gel, but intervening lanes were removed. (C) To determine if the two intermediate complexes (indicated with arrows) contain the same relative amount of H2A-H2B, FACT was mixed with an even larger excess of H2A-H2B (160 nM and 240 nM) before adding tetrasome.

Only the faster-migrating intermediate complex is formed at 60 nM H2A-H2B (Figure 4A, left panel lanes 5, 6). At high H2A-H2B concentrations (240 nM), the majority of the material was in the slower migrating species (Figure 4C, lane 4). Importantly, with 147 bp DNA, either intermediate complex only formed at > 20-fold excess of FACT and H2A-H2B over (H3-H4)_2_ tetrasome.

### DNA displaces bound FACT from the complex, and facilitates H2A-H2B deposition to assemble nucleosomes

To test whether DNA beyond the ~80 bp organized by the (H3-H4)_2_ tetramer contributes to the intermediate complex, we attempted to assemble it onto 79 bp DNA. The (H3-H4)_2_ tetrasome was first reconstituted with 79 bp DNA fragment, and then combined with FACT●H2A-H2B (Figure 5A). As shown before with longer DNA, FACT does not bind to the (H3-H4)_2_ tetrasome in the absence of H2A-H2B dimer (lane 3). Similarly, H2A-H2B does not bind properly to (H3-H4)_2_ tetrasome in the absence of FACT, but rather forms aggregates that do not enter the gel (lane 2). Thus, with 79 bp DNA, the intermediate complexes form only with tetrasomes and FACT-H2A-H2B, as observed previously with 147 bp DNA. However, no nucleosomes are formed, and unlike 147 bp DNA which only assembles into intermediate complexes at very high FACT and H2A-H2B concentrations, the intermediate complex with 79 bp DNA was observed at equimolar FACT to (H3-H4)_2_ to H2A-HB ratios.

**Figure 5.**
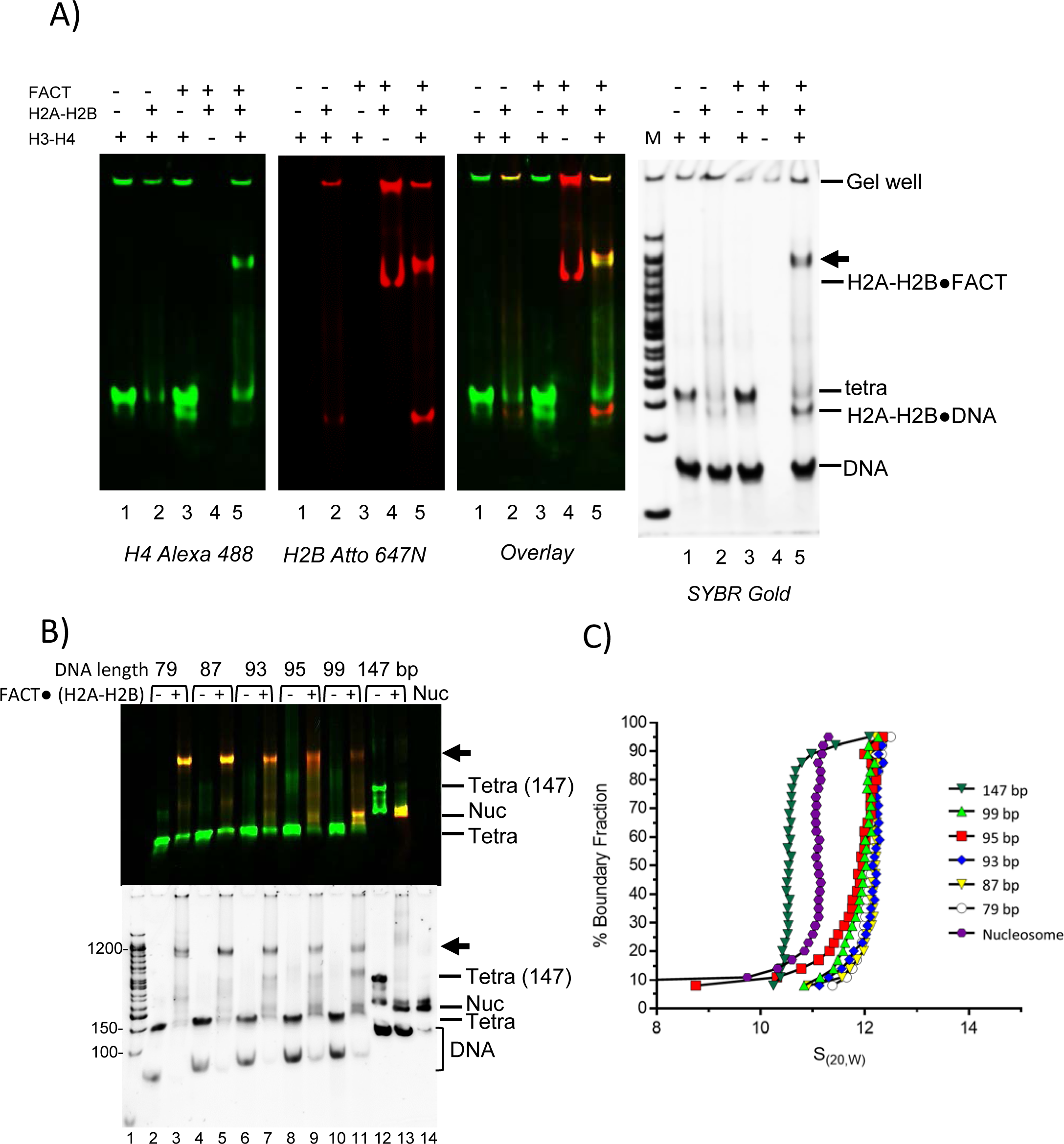
DNA competes FACT away from H2A-H2B, and facilitates formation of hexasomes/nucleosomes. (A) The intermediate complex is also formed with 79 bp DNA. FACT was pre-incubated with H2A-H2B for 10 min at room temperature, and tetrasome (pre-assembled with (H3-H4)2 and 79 bp DNA) was then incubated for 30 min at room temperature. In the final reaction, all components are at ~400 nM. The 5% native gel was visualized by SYBR Gold staining, or through H4 (Alexa 488) or H2B (Atto 647). (B) DNA competes FACT off H2A-H2B, and facilitates H2A-H2B deposition onto tetrasomes. FACT was pre-mixed with H2A-H2B, and tetrasome with different DNA length (79-147 bp) was added. In the final reaction, Alexa 488 labeled tetrasome concentration is ~500 nM. FACT is at 500 nM, and Atto 647N labeled H2A-H2B is ~1000 nM. The 5% native gel was visualized by H4 (Alexa 488, green) or H2B (Atto 647; red). The top panel shows an overlay of scans at both wavelengths. Bottom panel: SYBR Gold. (C) Complexes formed with different length DNA fragments, as in (B), were analyzed by SV-AUC (FDS at 488 nm) to monitor fluorescently labeled H4. S-values are listed in Table 1.

To better define the effect of DNA length in intermediate complex formation and nucleosome assembly, DNA of different lengths (79, 87, 93, 95, 99 and 147 bp) was tested. FACT, (H3-H4)_2_ tetrasome, and H2A-H2B were mixed at a 1:1:2 ratio. Only shorter DNA (79, 87, and 93 bp) form near-homogenous intermediate complexes (Figure 5B, lanes 3 and 5). With longer DNA (95 and 99 bp), the intermediate complex still forms, but the main products are partially assembled nucleosomes that contain H2A-H2B, H3-H4 and DNA, but no FACT. When the DNA is 147 bp long, almost no intermediate complex is formed, and the assembly product is mostly nucleosome (Figure 5B).

We analyzed these reactions by AUC-FDS at 488 nm, where only complexes containing fluorescently labeled H4 are visible (Figure 5C; S-values listed in Table 1). The intermediate complexes on either 79 bp (12.35S) or 87 bp (12.41S) DNA are quite homogenous in size, and only small amounts of a minor species with lower sedimentation values were observed. For complexes formed with 93, 95 and 99 bp DNA, the major species are also likely the FACT-bound intermediate complex, while the other components probably represent nucleosomes or their assembly intermediates in the absence of FACT. For the intermediate complexes with 147 bp DNA, the major species sediments as a nucleosome, with some hexasome and tetrasome, and little to no intermediate complex, consistent with what was observed by native PAGE (Figure 5B). Together, these experiments demonstrate that when free DNA extends from the tetrasome, it can compete FACT from H2A-H2B and dislodge it from the complex. This effectively results in H2A-H2B deposition onto (H3-H4)_2_ tetrasomes to form hexasomes, and ultimately nucleosomes. Thus, the intermediate complex is quite stable when the DNA is of limited length and unable to compete with FACT. In contrast, at least a 20-fold molar excess of FACT over (H3-H4)_2_ tetrasome is needed to ‘win’ the competition with DNA and to form the intermediate complex with 147 bp DNA. Importantly, our data demonstrate that we indeed observe intermediates states of FACT in the act of assembling nucleosomes or stabilizing partially disassembled nucleosomes.

### H2A-H2B interacts with both FACT and H3-H4 in the intermediate complex

Having established that H2A-H2B is required for the intermediate complex, we tested whether direct interactions between H2A-H2B and FACT are required. FACT ΔCTD (Spt16_1-932_) was previously shown to be deficient for H2A-H2B binding (Winkler et al., 2011). AUC-FDS confirmed that this complex does not bind to H2A-H2B, since no change in sedimentation was observed upon mixing 100 nM Alexa 488 labeled H2A-H2B with 200 nM FACT ΔCTD (Figure S5), and no intermediate complex was formed on a 79 bp DNA fragment (Figure 6A, lane 5). Instead, aggregates are formed similar to those obtained in the absence of FACT.

**Figure 6.**
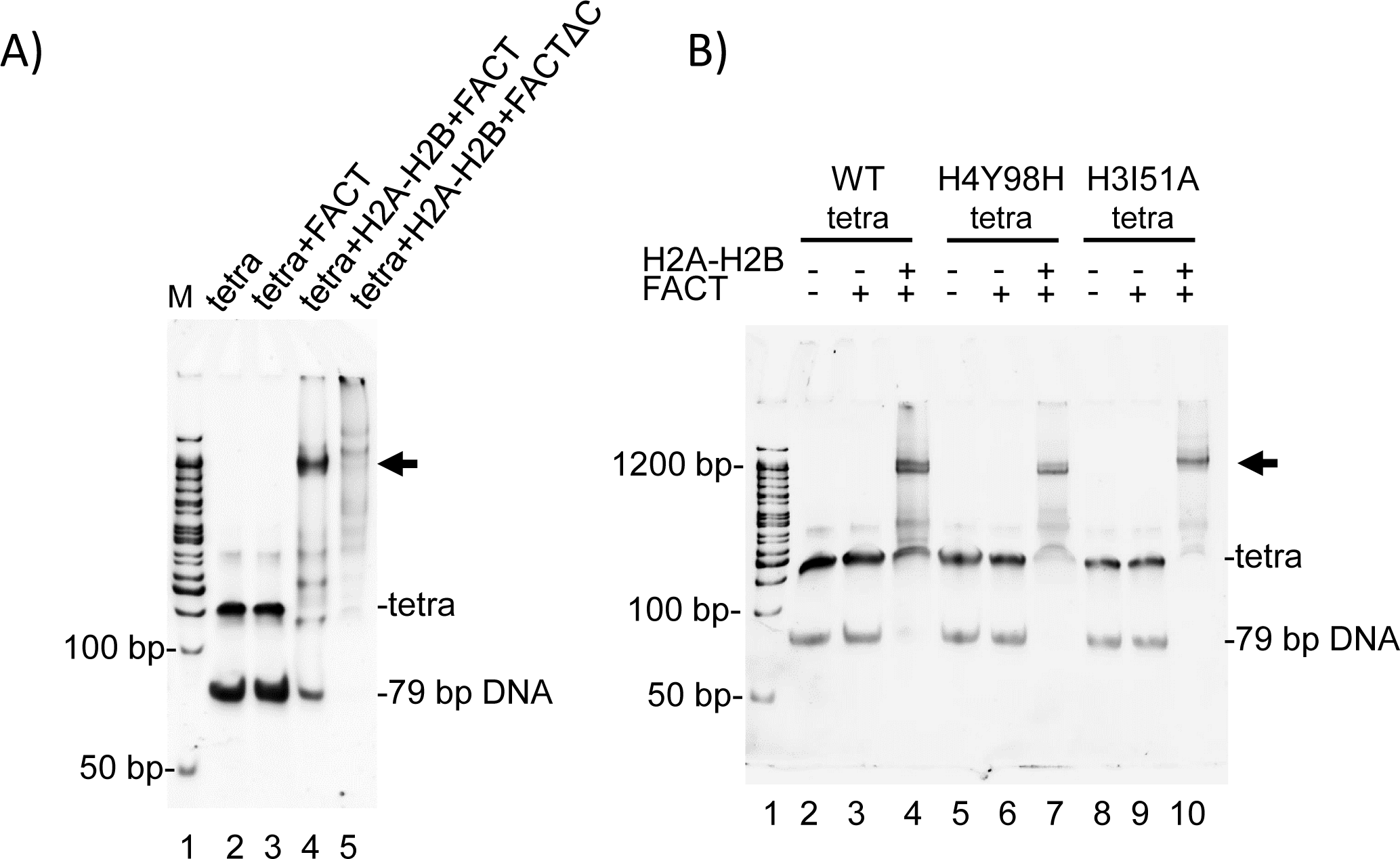
FACT●H2A-H2B interactions are required for the intermediate complex. (A) 500 nM FACT or FACT ΔC was pre-mixed with 1000 nM H2A-H2B, then tetrasome with 79bp DNA (500 nM) was added. Intermediate complex is indicated by an arrow. (B) FACT●H2A-H2B was incubated with WT or mutant (Y98H or I51A) tetrasome, reconstituted with 79 bp DNA. Concentrations were as in (A).

Next, we queried the requirement for direct H2A-H2B and H3-H4 interactions, by testing mutant histones H3I51A and H4Y98H (Figure 1C) on a 79 bp DNA fragment. H3I51A does not significantly impede the formation of intermediate complex, while intermediate complex formation with H4Y98H was moderately compromised (Figure 6B). This indicates that H2A-H2B also interacts with H3-H4 in the intermediate complex, as it does in the histone hexamer-FACT complex (Figure 1D).

### FACT facilitates transcription by RNA polymerase II

Finally, we wanted to re-investigate the effect of FACT on transcription by using a recombinant in vitro transcription system. In previous reports, FACT was shown to have only a moderate effect on the amount of full length RNAPII transcript (Kuryan et al., 2012). In light of our finding that FACT binds to hexasomes, we wanted to test if FACT affects yeast RNAPII pause sites. To this end, nucleosomes were reconstituted on the templates shown in Figure 7A. Nucleosomes were kept at 0.5 nM, and FACT was added at the indicated amounts. Transcription was initiated by the addition of NTP, and RNA transcripts were analyzed on a sequencing gel (Figure 7B). Nucleosomes preclude formation of full-length transcript (compare Figure 7C, lanes 2, 3), and this repression was relieved upon addition of the ATP-dependent chromatin remodeler RSC and the histone chaperone Nap1 (lane 5). FACT on its own has only moderate effects on the amount of full length transcript, whereas in the presence of RSC it stimulates transcription in a non-linear dose-dependent manner, consistent with previous results (Kuryan et al., 2012, Orphanides et al., 1999). Notably, two shorter transcripts indicative of polymerase pause sites were observed in the presence of FACT, one at ~85-bp, or 35 bp into the nucleosome, and one at ~105 bp, or 55 bp into the nucleosome (Figure 7C, lane 6-10, and Figure 1C). The first site coincides with the interaction of DNA with the H2B L1 loop, and is thus consistent with polymerase passage through DNA without being able to progress through the H4-H2B four-helix bundle. The second pause site coincides with the H3-H4 α1α1 interface, and could stem from the interaction of FACT with a hexasome (i.e. the intermediate complex described above). The abundance of this shorter RNA increases with increasing FACT concentration (and decreasing full length transcript). In the presence of RSC, no polymerase pausing is observed at this location.

**Figure 7.**
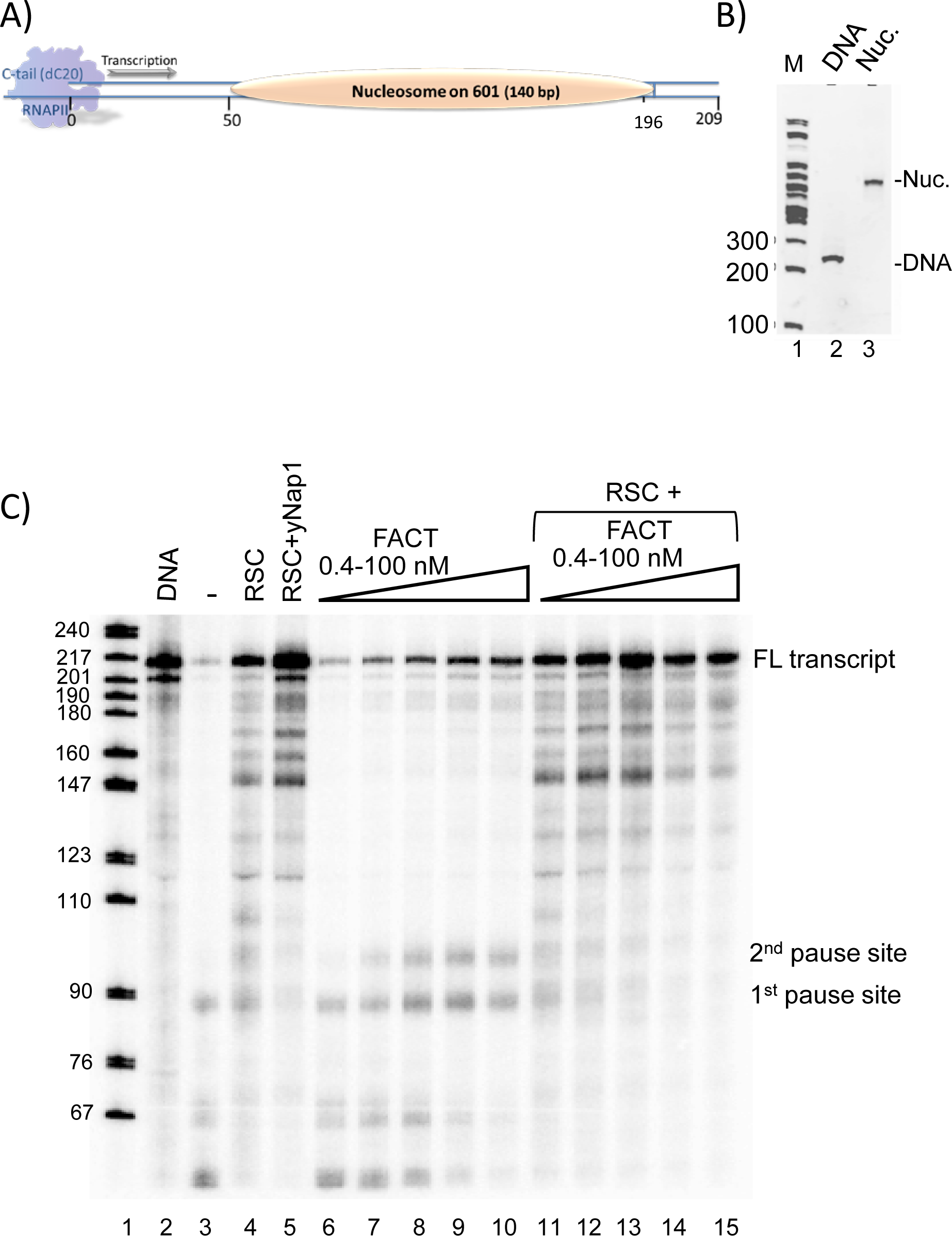
FACT affects RNAPII-mediated transcription through a nucleosome. (A) Schematic of the transcription template. (B) Nucleosomes were reconstituted on the transcription template shown in (A), and analyzed by native PAGE. Note the absence of free DNA. (C) The transcription output was analyzed on a 6.5% acrylamide sequencing gel, and visualized by radioactivity (P32). Pause sites are also indicated in Fig. 1C

## Discussion

The mechanism by which FACT, an essential and conserved histone chaperone complex, facilitates the processes of gene transcription, DNA replication and DNA repair in the context of chromatin is unknown (Keller et al., 2002, Schlesinger et al., 2000, Wittmeyer et al., 1997). Our results with purified FACT and nucleosomes / histones provide mechanistic insight into how hFACT stabilizes intermediate states of the nucleosome, seemingly promoting both nucleosome disassembly and assembly, and ultimately regulating access to nucleosomal DNA.

FACT alone does not bind to or disassemble fully assembled nucleosomes, consistent with recent data (Tsunaka et al., 2016), but contradicting published work with HeLa nucleosomes in pull-down assays (Belotserkovskaya et al., 2003). HeLa nucleosomes carry an abundance of post-translational modifications and likely exist in a mixture of intact and partially destabilized nucleosomes, which are further destabilized with an excess of FACT. In our own published work using FRET-based assays, we have determined that FACT interacts with nucleosomes (Winkler et al., 2011). The observed FRET changes might be attributed to interactions of FACT with free labeled H2A-H2B dimer, or with hexasomes that might exist at these very low nucleosome concentrations. Analytical ultracentrifugation experiments provide first-principle information on the behavior of macromolecules in solution, and allowed us to determine the inability of FACT to bind to intact nucleosomes. FACT is also unable to disassemble nucleosomes, and likely requires destabilizing activities such as RNAPII or ATP-dependent chromatin remodeling factors for its action.

FACT is particularly relevant during gene transcription, where the requirement for *de novo* nucleosome assembly is not as stringent as during replication or DNA repair. Especially at slower transcription rates, tetrasomes or hexasomes survive the passage of RNAPII (Dion et al., 2007, Jamai et al., 2007, Thiriet et al., 2006). Indeed, we found that FACT is only moderately effective in mediating the deposition of (H3-H4)_2_ tetramer onto DNA. FACT can deposit an H2A-H2B dimer onto a tetramer to form a hexasome, whereas the addition of the second H2A-H2B dimer is not as favorable and requires a large excess of FACT.

FACT, when bound to a histone H2A-H2B dimer, assembles into a stable intermediate complex with (H3-H4)_2_ bound to DNA (a tetrasome). This complex differs from the complex with histones in the absence of DNA described previously (Tsunaka et al., 2016) and confirmed here. Because 79 bp of DNA, the minimal length to wrap the entire (H3-H4)_2_ tetramer, are sufficient to form this complex, a significant portion of the 147 bp DNA that would be tightly bound in the context of a nucleosome is freely accessible. We suggest a model where free DNA (when not otherwise engaged with a transcribing RNAPII, or bound by transcriptional regulators), can compete with FACT for H2A-H2B, resulting in FACT displacement and the formation of a hexasome, which appears to be the predominant end-product of FACT-mediated nucleosome assembly *in vitro* (Figure. 8). This explains why the intermediate complex is observed on longer DNA fragments only when a large excess of FACT is used, and implies that the interaction of the FACT-H2A-H2B complex with a hexasome to potentially deliver the second dimer is unfavorable. A stable complex between FACT and a hexasome, in which only ~80 bp of DNA are tightly bound, is also consistent with the idea that yeast FACT ‘reorganizes nucleosomes’ (Xin et al., 2009). A similar complex was observed with a nucleosome that had been reconstituted with two DNA fragments (33 and 112-bp), which significantly destabilizes the nucleosome and results in the loss of one H2A-H2B dimer, and possibly the short DNA fragment (Tsunaka et al., 2016).

**Figure 8.**
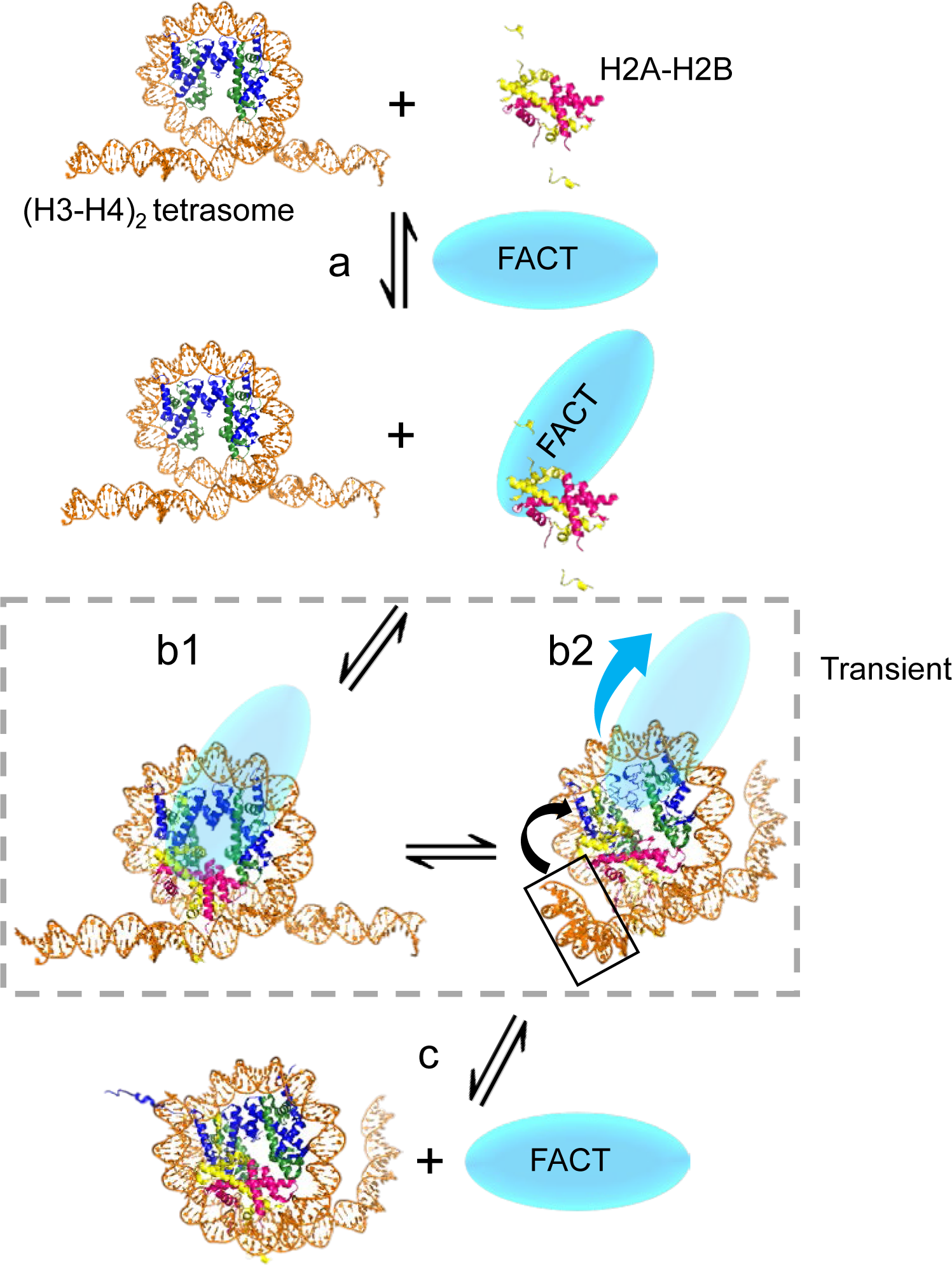
Model of FACT action in nucleosome/hexasome assembly. Step a. FACT binds to H2A-H2B dimer; Step b1. FACT chaperones H2A-H2B onto (H3-H4)_2_ tetrasome; Step b2. free DNA end competes FACT off H2A-H2B; Step c. Hexasome is formed.

H2A-H2B does not interact with H3-H4 in the absence of DNA at physiological conditions, but is able to do so when pre-bound to FACT. The interactions between these histones in the FACT complex are similar to the nucleosome. FACT SPT16 CTD interaction with H2A-H2B dimer is required to form a complex with a hexasome. The contributions of other functional domains of FACT are still unknown, but some are likely to also participate. For example, recent studies with an artificially tethered HMGB domain highlight its importance for yeast FACT function (McCullough et al., 2018).

Our results suggest a mechanism by which FACT facilitates gene transcription through a nucleosome. Shorter transcripts resulting from RNAPII pausing are uniquely observed in the presence of FACT, independent of the chromatin remodeling factor RSC. Because the position of the nucleosome on 601 DNA is known with high accuracy, these pause sites give insight into where FACT might interact with nucleosomal DNA during transcription (Figure 1C). The concept that FACT tethers partially assembled nucleosomes is consistent with the finding that histone turnover rates are enhanced in yeast when FACT activity is compromised (Jamai et al., 2009). During gene transcription, H2A-H2B is much more dynamic than H3-H4 (Jamai et al., 2007). FACT might capture an H2A-H2B dimer that had been displaced from the nucleosome by the advancing RNA polymerase, and re-instate it on the tetrasome by forming a ternary complex such as the one we describe here. In this model, FACT is displaced by DNA, once that DNA is no longer engaged with the polymerase (Figure 8). FACT or another H2A-H2B chaperone could then facilitate deposition of the second H2A-H2B onto a hexasome (without forming a ternary complex) to complete nucleosome reassembly. Similarly, FACT could temporarily stabilize a partial nucleosome complex consisting of one H2A-H2B dimer (whose DNA is engaged with the polymerase) and one DNA-bound (H3-H4)_2_ tetramer. Our model explains the dose-response curve of FACT on transcript lengths, because at very high FACT concentrations it can no longer be displaced and inhibits transcription.

## Materials and Methods

### Reagents

Recombinant histone genes (sequences derived from either *Xenopus laevis* or *Homo sapiens*) were expressed and purified as described previously (Luger et al., 1999). H4T71C mutant was used for labeling of H3-H4 with Alexa 488, while H2BT112C was used to label H2A-H2B with Atto 647N. 147 bp or 207 bp 601 DNA was prepared as described, and nucleosomes with 147 bp or 207 bp DNA were reconstituted by salt dialysis (Dyer et al., 2004).

### FACT expression and purification

The purification of human FACT (full length or FACT with Spt16_1-934_; ΔCTD) was adapted from published work with minor changes (Winkler et al., 2011). FACT complexes were purified over a 5 ml prepacked HisTrap HP column, followed by purification over a 5 ml prepacked HiTrap Q HP column. The final step was a Superdex 200 10/300 size exclusion column in 300 mM NaCl, 20 mM Tris pH 8.0, 5% glycerol, 0.01% CHAPS, 0.01% octyl glucoside and 1 mM TCEP. FACT was stored in 150 mM NaCl, 20 mM Tris pH 8.0, 10% glycerol, 0.01% CHAPS, 0.01% octyl glucoside and 1 mM TCEP (buffer A). CHAPS and octyl glucoside help avoid nonspecific protein-protein interaction. All columns were purchased from GE Healthcare.

### Sedimentation velocity analytical ultracentrifugation (SV-AUC)

To determine the stoichiometry of the FACT-(H2A-H2B) complex, Sedimentation Velocity Analytical Ultracentrifugation (SV-AUC) with absorbance optics using intensity mode (collected at 280 nm) was performed in buffer containing 150 mM NaCl, 20 mM Tris pH 8.0 and 1 mM TCEP. 1.8 µM FACT was incubated with the indicated amount of H2A-H2B at room temperature for 10 min, then spun at 30-35,000 rpm and 20 °C in a Beckman XL-A ultracentrifuge, using an An60Ti rotor. Partial specific volumes of the samples were determined using UltraScan 3 version 2.0. Time invariant and radially invariant noise was subtracted using 2-dimensional-spectrum analysis (2DSA) followed by genetic algorithm refinement and Monte Carlo analysis to resolve the sedimentation coefficients (s, converted to Svedberg units (S) by multiplying by 10^−13^ and corrected for water at 20 ° Celsius) and frictional ratios (f/f_0_) of significant species in each sample, from which the apparent molecular weights were extracted (Brookes et al., 2010); (Cao et al., 2008) (Brookes et al., 2010). Sedimentation coefficient distributions G(s) were obtained with enhanced van Holde-Weischet analysis, plotted with sedimentation coefficients (s) converted to Svedbergs (Demeler et al., 2004). Analyses were performed using Ultrascan 3 version 2.0, and distributions were plotted using Graphpad Prism.

To determine if FACT binds nucleosomes, 150 nM nucleosome was incubated with 600 nM FACT at room temperature for 10 min, then spun using an An50Ti or An60Ti rotor (Beckman Coulter) at 32,000 rpm, 20 °C in a Beckman XL-A ultracentrifuge using absorbance optics at 280 nm. Data analysis was performed as described above.

To determine if FACT binds a histone hexamer, SV-AUC experiments were performed in a Beckman XL-A ultracentrifuge equipped with an Aviv fluorescence detection system (FDS), using an An60Ti rotor (Beckman Coulter) with standard epon 2-channel centerpiece cells. Alexa 488 labeled *Xenopus laevis* histones (maleimide linked at either H4 T71C or H2B T112C) were fixed at either 100 or 200 nM and incubated with FACT. Sedimentation was then monitored with fluorescence optics (excitation 488 nm, emission >505 nm) at 20 °C using speeds of 40 or 45,000 rpm.

To determine the effect of DNA length on FACT●H2A-H2B●(H3-H4)_2_ complex formation, SV-AUC experiments were performed in a Beckman XL-A ultracentrifuge with Aviv FDS, using an An50Ti rotor with standard epon 2-channel centerpiece cells. *Homo sapiens* histones were labeled at H4 T71C via maleimide linkage with Alexa 488 and were reconstituted with H3 and DNA with the indicated lengths (79, 87, 93, 95, 99 or 147 bp) to form (H3-H4)_2_ tetrasomes by salt reconstitution. Labeled tetrasomes were adjusted to 350 nM (in buffer containing 20 mM Tris HCl pH 7.5, 150 mM NaCl, 1mM EDTA), and combined with equimolar amounts of pre-incubated FACT●H2AB. Sedimentation was then monitored using fluorescence optics at 20 °C and 30,000 rpm. Ultrascan 3 version 4.0 was used for all AUC-FDS analysis, and distributions plotted using Graphpad Prism.

All data analysis was performed on the UltraScan LIMS cluster at the Bioinformatics Core Facility at the University of Texas Health Science Center at San Antonio and the Lonestar cluster at the Texas Advanced Computing Center supported by NSF Teragrid Grant #MCB070038.

### Nucleosome assembly/disassembly assay

To test if FACT disassembles nucleosomes, FACT (5 nM to 5 µM) was titrated into nucleosomes previously assembled onto 147 bp ‘601’ DNA (10 nM), and incubated in buffer A at room temperature for 1 h, then analyzed by 5% PAGE, run at 150 V, 4 ˚C for 60 min.

To determine if FACT facilitates tetrasome assembly, FACT (15-240 nM) was titrated into refolded (H3-H4)_2_ tetramer (30 nM), and incubated in buffer A at room temperature for 10 min. DNA (30 nM) was added, and tetrasome formation was analyzed by native PAGE as described above.

To determine if FACT facilitates H2A-H2B deposition onto tetrasome, Atto 647N labeled H2A-H2B dimer (30 nM) was mixed with varying amounts of FACT, and incubated at room temperature for 10 min. 60 nM tetrasome was added and incubated at room temperature for another 30 min. The reaction was performed in buffer A and analyzed as described above.

To test whether FACT facilitates H2A-H2B deposition onto hexasomes, under-assembled nucleosomes were reconstituted using an optimal ratio of H3-H4 to DNA and sub-saturating amounts of H2A-H2B. FACT (10 nM to 1280 nM) was then titrated into 20 nM under-assembled nucleosome, and analyzed as described above.

To investigate if FACT facilitates nucleosome assembly, Alexa 488 labeled H3-H4 (500 nM) and Atto 647N labeled H2A-H2B dimer (500 nM) were mixed with or without FACT (1 µM), and incubated at room temperature for 10 min. 207 bp 601 DNA (250 nM) was added, incubated at room temperature for 30 min, and analyzed by native PAGE.

To determine if FACT assembles super-shifted complexes with tetrasome with either 147 or 79 bp DNA, 850 nM FACT was mixed with increasing amounts of H2A-H2B (160 nM and 240 nM), and then 40 nM tetrasome with either 147 or 79-bp DNA was added.

### Micrococcal nuclease digestion assay

3.2 µM FACT was mixed with 400 nM H3-H4 and 400 nM H2A-H2B dimer, and incubated at room temperature for 10 min. 100 nM 147bp DNA was then added, and incubated at room temperature for 30 min in buffer A. Increasing amounts of MNase (100 U, 200 U, 400 U) was added to the Anti-FLAG purified supershifted complexes, and incubated at 37 °C for 10 min. The reaction was quenched by adding 5 µl 0.5 M EDTA. 621 bp DNA was added as a recovery standard. DNA fragments were purified through a MiniElute PCR Purification Kit (QIAGEN). DNA fragments were quantified by 2100 Bioanalyzer (Agilent), and analyzed by 2100 Expert software. To determine the stability of nucleosome assembled by FACT, MNase digestions for FACT-assembled nucleosomes were performed and analyzed as described above.

### In vitro transcription assay

The in vitro transcription assay was adapted from published work (Kuryan et al., 2012). 141 bp 601 DNA, flanked by 50 bp polylinker DNA and 20-bp plasmid DNA were joined to a single-stranded C tail as a binding site for RNAPII (sequence given below). Mononucleosomes were reconstituted by salt dialysis; special care was taken to reach a correct assembly ratio without free DNA. All transcription reactions were performed in 25 mM HEPES pH 7.5, 10 mM MgCl_2_, 2.5 mM KCl, 10% glycerol, 1 mM DTT, and 250 ng/µl BSA. Nucleosome concentration was kept at 0.5 nM (free DNA at 0.1 nM). FACT was titrated from 0.4 nM to 100 nM, and transcription was initiated by adding dNTP. RNA transcripts were analyzed on a 6.5% acrylamide sequencing gel. RSC and RNA pol II were prepared as previously described (Izban et al., 1991).

### C-tail sequence

~~~
CCCCCCCCCCCCCCCCCCCCTGTGGGCCCTTCTTTTTCGTTTGGCGTCTCTAGACACCCGGGAAGAAAAAGCAAACCGCAGA
~~~

Ligated to 180 bp-601 sequence (only bottom strand shown, BamH1 and EcoRV sites underlined)

~~~
**GATCC**ATGCACA<u>**GGATGTA**</u>TATATCTGACACGTGCCTGGAGACTAGGGAGTAATCCCCTTGGCGGTTAAAACGCGGG
~~~

~~~
GGACAGCGCGTACGTGCGTTTAAGCGGTGCTAGAGCTGTCTACGACCAATTGAGCGGCCTCGGCACCGGGAT<u>**TCTCCA**</u>GGAATTCAAGCTTCCCGGGGGGGAT
~~~

## Acknowledgements

We thank Dr. Serge Bergeron for valuable suggestions regarding the nucleosome assembly assay. Supported by the Howard Hughes Medical Institute, by NIH-GM-067777, and by 1F31GM105363 to DDK

## Author Contributions

Tao Wang and Yang Liu: validation, investigation, formal analysis, methodology, writing and editing of manuscript; Daniel Krzizike and Garrett Edwards: AUC experiments; Hataichanok Scherman: in vitro transcription assay; Karolin Luger: concept and experimental design, manuscript preparation.

## Supplementary Figure legends

**Figure S1.**
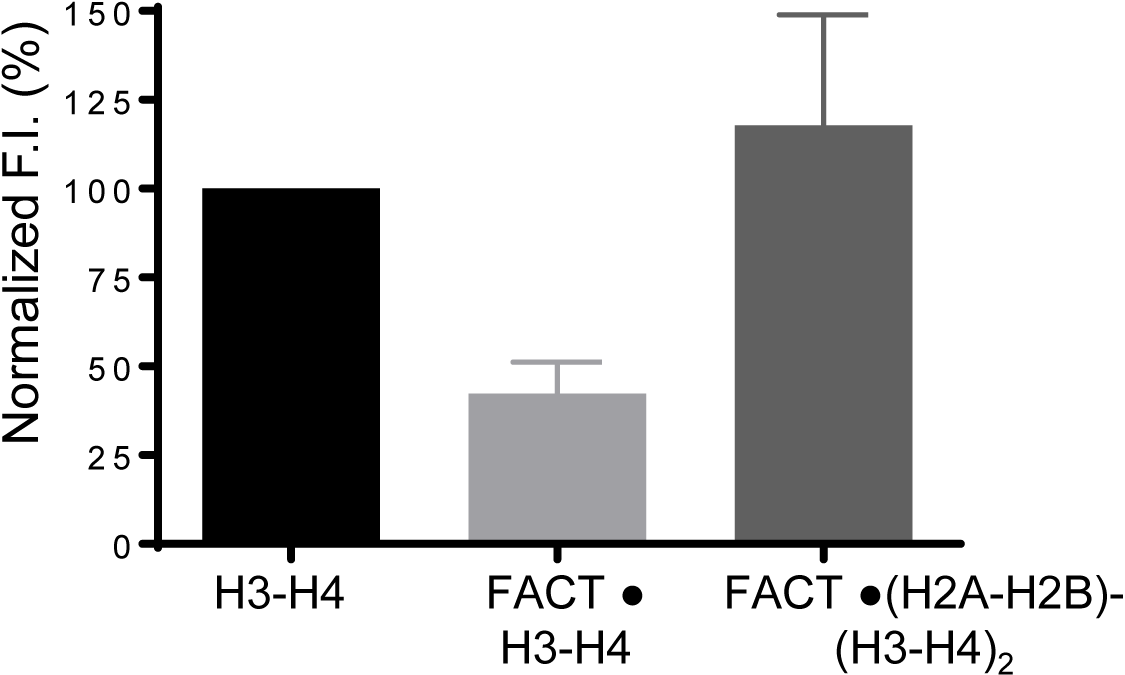
H2A-H2B prevents FACT from forming aggregates with H3-H4. We attempted to determine the S(20,W) value of the FACT-H3-H4 complex by using AUC-FDS. Even at H3-H4 concentration as low as 200 nM, FACT forms aggregates with H3-H4, causing ~50% signal loss. Upon addition of H2A-H2B, signal is regained, suggesting that H2A-H2B facilitates the formation of an ordered complex between FACT and H3-H4. The error bars represent SD from two biological replicates.

**Figure S2.**
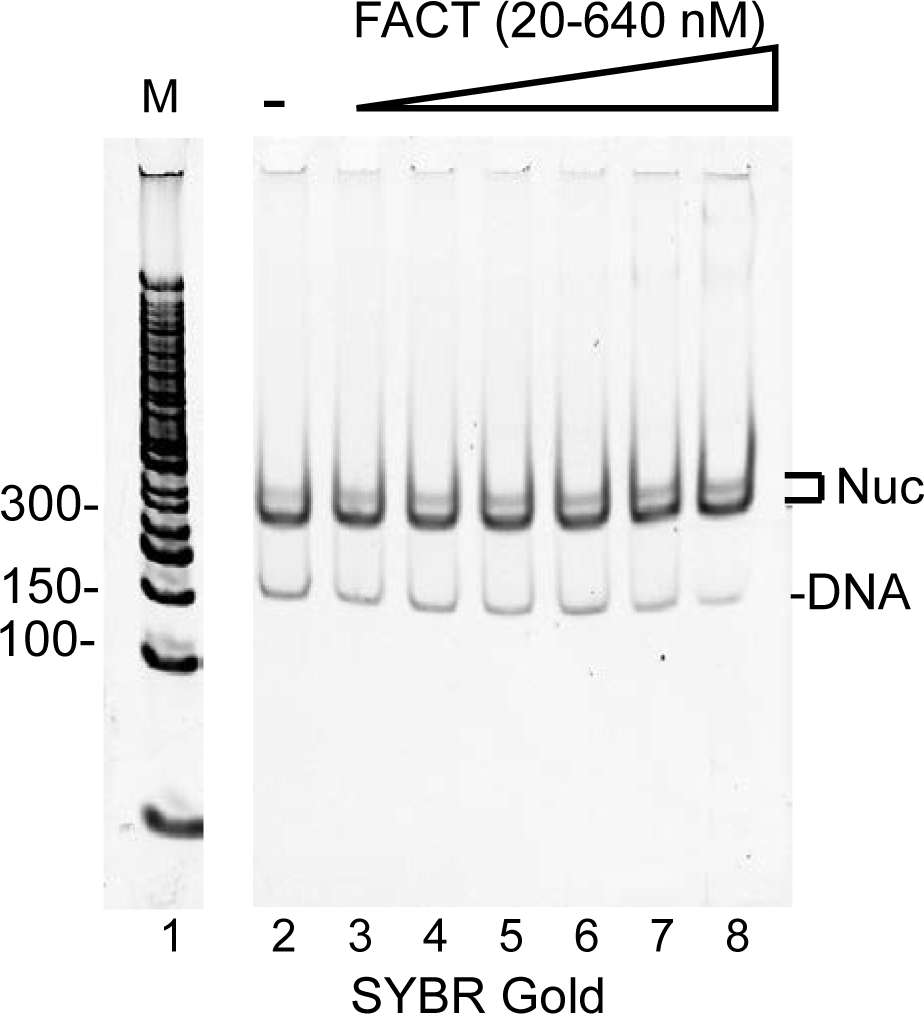
FACT does not disassemble nucleosomes assembled on α-satellite DNA. Nucleosomes were reconstituted with 147-bp α-satellite DNA and *Xenopus laevis* histones. Nucleosome concentration was kept at 20 nM. FACT was added at the indicated concentrations, and samples were incubated at 40 °C for 1 h. Samples were analyzed by 5% PAGE, and visualized by SYBR Gold. In absence of FACT, nucleosomes remain intact at 40 ˚C (lane 2). As FACT is titrated, the amount of nucleosome and free DNA is unchanged (lanes 3 to 8).

**Figure S3.**
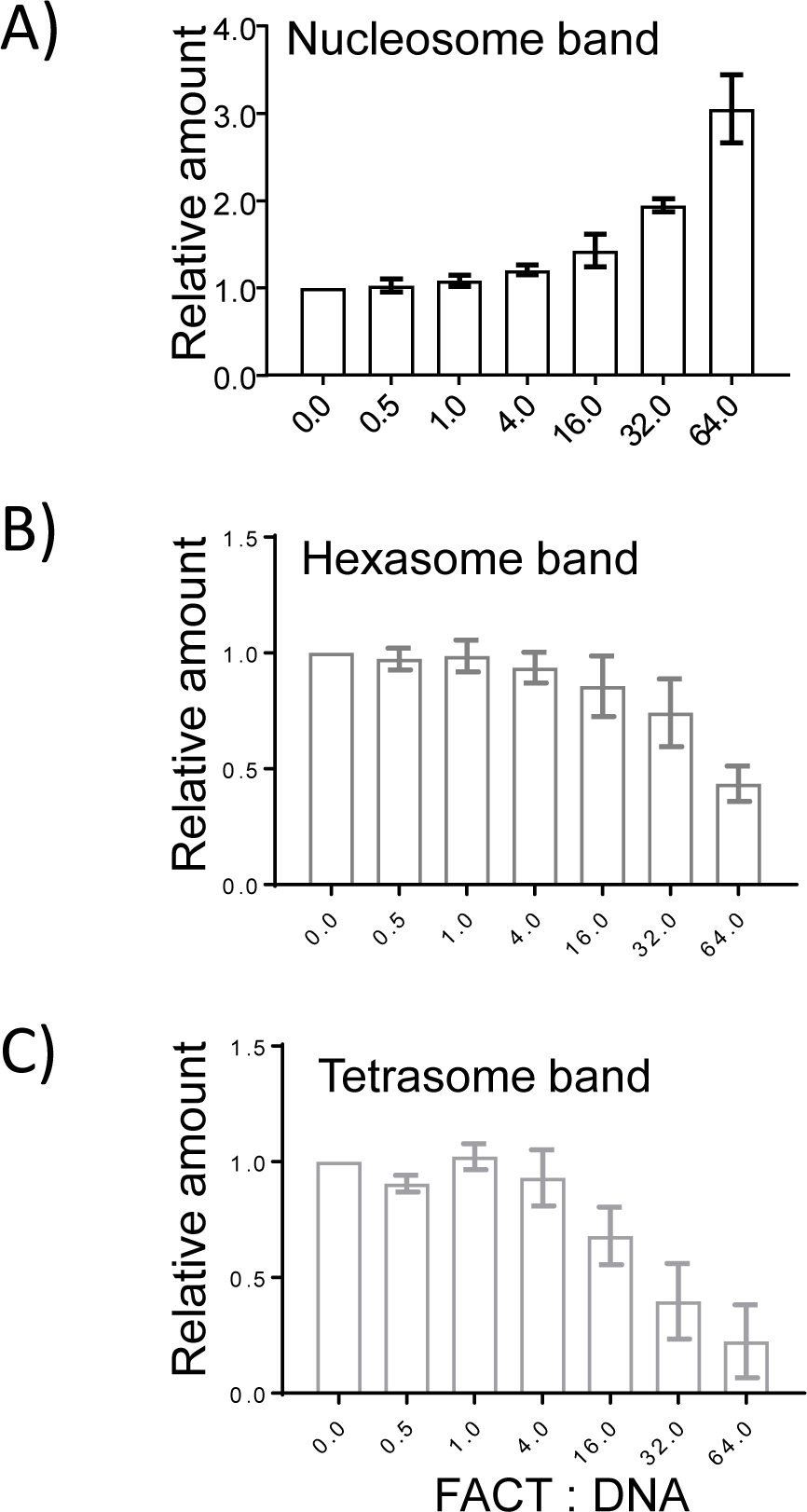
FACT facilitates the deposition of H2A-H2B onto a hexasome. The gel shown in Fig. 3C (and two others) were quantitated by ImageQuant™. The fluorescence signals from nucleosomal H4 (A), hexasomal H2B (B) and tetrasomal H4 (C) were quantified. The intensity of controls (nucleosome, hexasome or tetrasome; lane 3) was assumed as 1, and the intensity ratio between samples with FACT and the control was determined. The error bars represent SD from three biological replicates.

**Figure S4.**
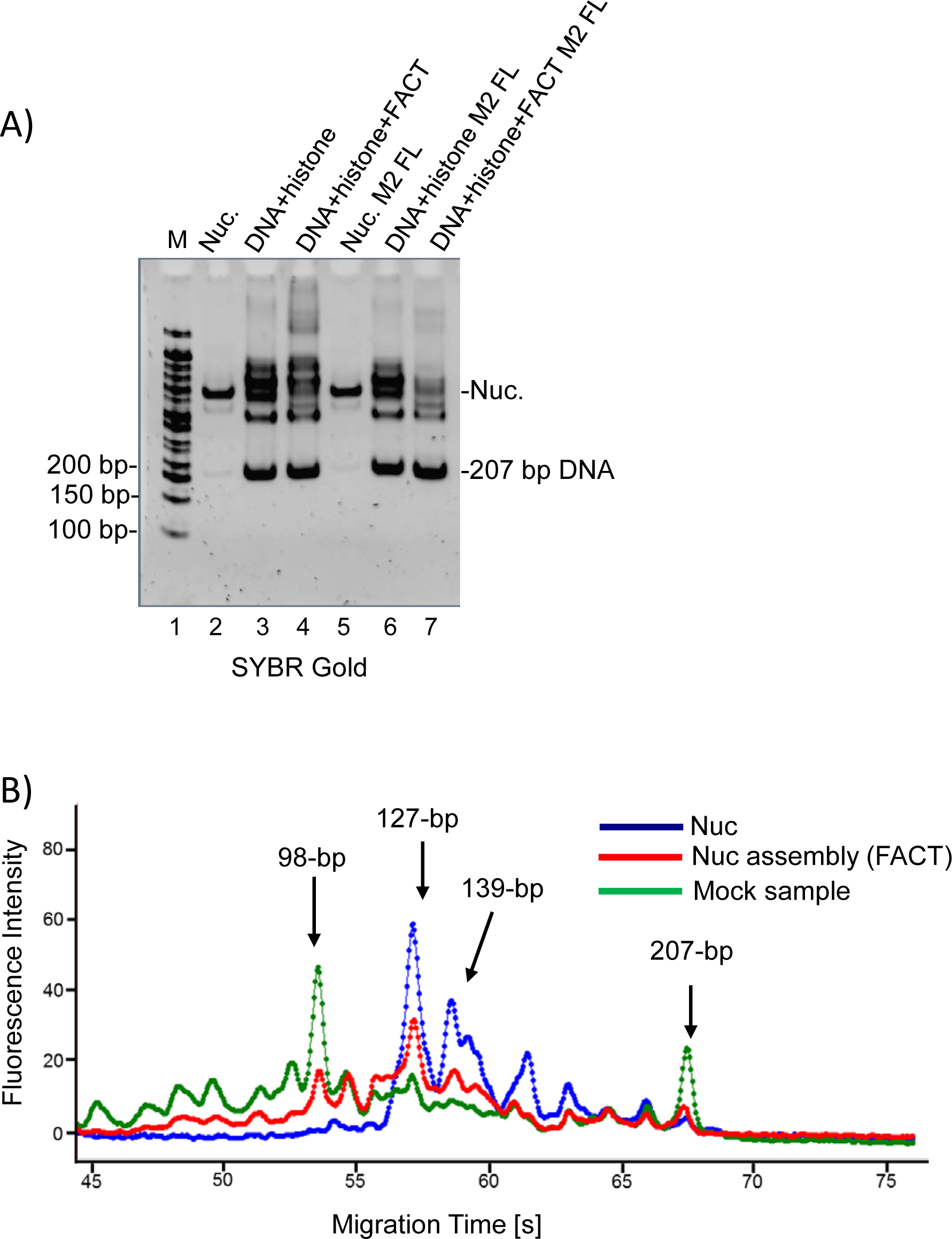
Nucleosomes assembled with FACT have similar MNase digestion pattern as salt-reconstituted nucleosomes. 1 µM FACT was mixed with 250 nM (H3-H4)2 tetramer and 500 nM H2A-H2B dimer, then 250 nM 207 bp 601 DNA was added. The supershifted FACT-containing complex was removed by M2 resin, and the flow through (containing various assembly products not bound by FACT) was collected for MNase digestion. Protected DNA fragments were quantified using a Bioanalyzer. Nucleosomes assembled by FACT (red trace) have similar MNase digestion pattern compared to salt-reconstituted nucleosome (blue line). A different pattern is observed in absence of FACT (mock sample, green line).

**Figure S5.**
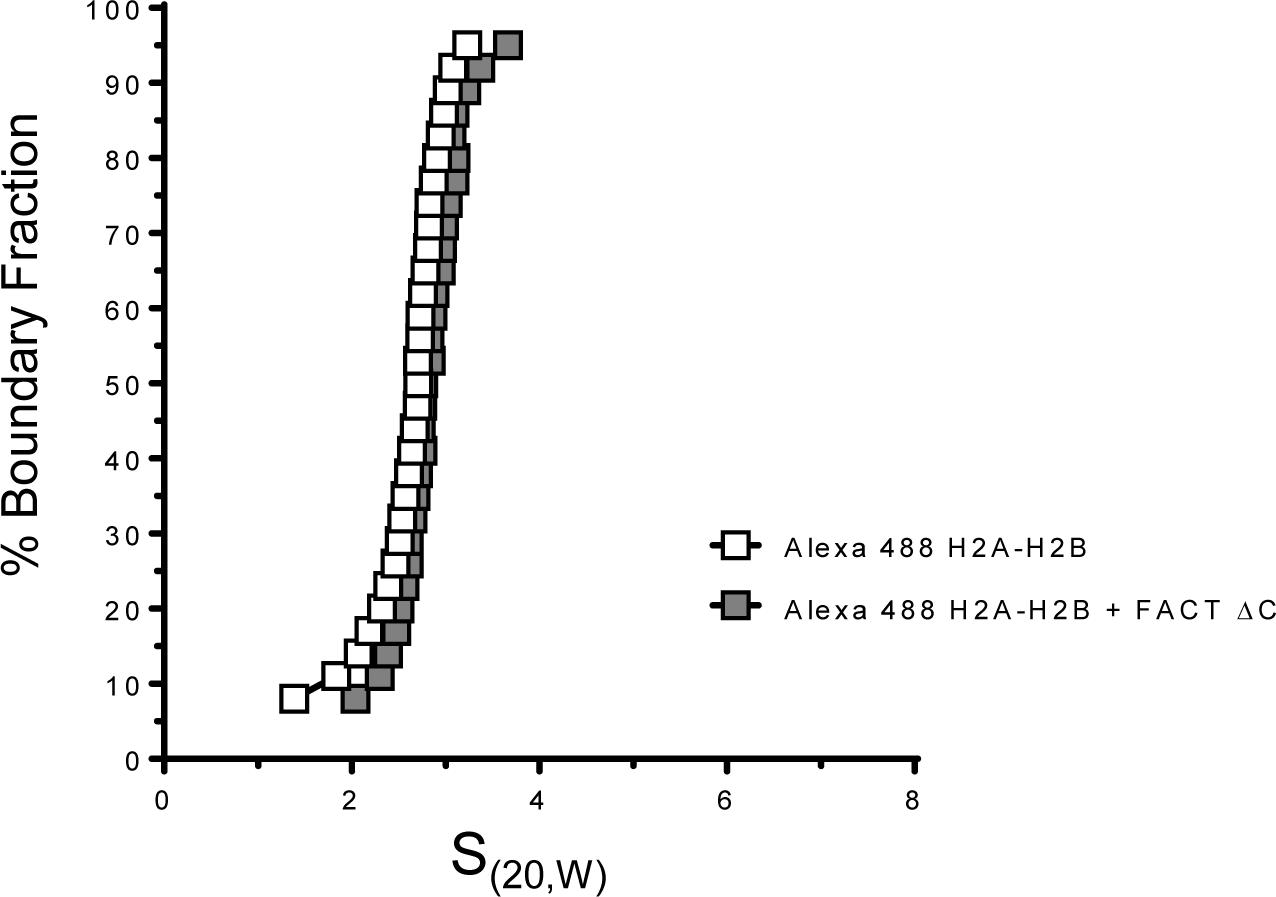
FACT ΔCTD does not bind to H2A-H2B with high affinity. 100 nM Alexa 488 labeled H2A-H2B was mixed with 200 nM FACT ΔC, and SV-AUC was performed by monitoring fluorescence using the FDS. Addition of FACT ΔC does not alter the S(_20,W_) value of H2A-H2B. The trace for Alexa 488 H2A-H2B is the same as the one shown in Fig. 1B.

**Table S1:** FACT facilitates nucleosome and hexasome assembly. 15 and 1500-bp DNA fragments are internal markers. MNase 400 U

| *Size (bp)* | *207 bp Nucleosome* | *207 bp DNA+histone*<br>*w/o FACT* | *207 bp DNA+histone*<br>*w/ FACT* |
| --- | --- | --- | --- |
|  | Molarity (nmol/l) | Molarity (nmol/l) | Molarity (nmol/l) |
| 15 | 424.4 | 424.4 | 424.4 |
| 98 | 0 | 16.4 | 0 |
| 106 | 0 | 0 | 11.6 |
| 115 | 0 | 0 | 9.9 |
| 126 | 34.7 | 0 | 19.5 |
| 139 | 6.9 | 0 | 0 |
| 1500 | 2.1 | 2.1 | 2.1 |

